# Frontoparietal Hub Connectivity Integrates Information from Multiple Sources

**DOI:** 10.64898/2026.04.09.717528

**Authors:** Stephanie C. Leach, Shannon E. Stokes, Jiefeng Jiang, Kai Hwang

## Abstract

Frontoparietal connector hubs are thought to support information integration across the brain, but this role has largely been inferred from static connectivity, leaving unclear how computational processes shape inter-regional connectivity during behavior. Here, we address this question using a model-based functional connectivity approach in human fMRI data. Thirty-Eight participants (males and females) performed a task requiring the integration of sensory evidence with an internally maintained state belief to guide behavior. We developed a computational model that combines these information sources into an integrated representation and generates distinct variables at successive stages of integration: uncertainty before choice (entropy), the inferred task representation guiding action (task belief), and feedback-driven updating (task inference error). We then tested how these variables modulate the connectivity of frontoparietal connector hubs. Entropy increased coupling between hubs and regions encoding task-relevant inputs and outputs during cue processing, suggesting enhanced communication under uncertainty. During task selection, task belief selectively modulated hub connectivity with motor regions according to the selected task. During feedback, task inference errors increased coupling with regions supporting task-relevant inputs and internal state, while reducing coupling with motor regions, consistent with updating internal representations. Together, our findings show that frontoparietal connector hubs implement integrative control by using an integrated representation to generate distinct computational signals that selectively and dynamically reconfigure inter-regional communication.

**Significance:** Flexible behavior depends on combining different kinds of information, but how the brain coordinates this integration remains unclear. The frontoparietal cortex is well positioned to support this process because it is broadly connected with many other systems. Here, we combined a computational model with functional MRI to test how integrating information changes patterns of functional connectivity. We find that a common set of signals is associated with dissociable changes in how frontoparietal regions couple with systems involved in perception, action, and internal updating. These findings reveal that integration generates multiple control signals that dynamically reconfigure brain-wide interactions to support goal-directed behavior.

## 1. Introduction

Integration is a central concept in neural and cognitive sciences. Integration combines multiple factors to jointly influence a process or outcome, and is essential for coordinating systems composed of subsystems (Damasio, 1989; Fuster, 2015). Everyday behavior depends on integration. For example, driving in unpredictable traffic requires combining multiple sources of information, including environmental sensory inputs (e.g., position/speed of other vehicles), working memory content (e.g., recent navigation instructions), and contextual knowledge (e.g., anticipated traffic conditions). To guide behavior, multiple streams of information are integrated to support goal-directed, adaptive behavior.

The need for integration has long been emphasized in theories of brain network organization (Mesulam, 1990; Tononi et al., 1994). One solution is to develop domain-general brain regions with extensive connectivity to distributed subsystems (Sporns and Betzel, 2016; van den Heuvel and Sporns, 2013). Extensive work indicates that regions in the frontoparietal cortices are extensively connected to diverse brain networks and function as “connector hubs” (Bertolero et al., 2015; Power et al., 2013). Because connector hubs link otherwise segregated systems, they are well positioned to integrate diverse sources of information to bias processing in task-relevant regions (Bertolero et al., 2018; Cole et al., 2013; Goldman-Rakic, 1988). Frontoparietal hubs are therefore strong candidates for supporting integrative control.

Although frontoparietal hubs are topologically positioned to integrate information, the computational signals that drive their integrative function remain unclear. In recent work, we developed a computational model describing how multiple task-relevant information sources converge in frontoparietal cortex and are combined into an integrated task representation (Leach et al., 2025). In the task, sensory evidence and contextual state information are combined to determine the action. The model formalizes this integration as a joint probability distribution over tasks—an integrated representation that encodes the subject’s belief about which task to perform.

Following feedback, state and task beliefs are updated to improve subsequent performance. This model generates several model-derived variables corresponding to different stages of the integration process. Entropy quantifies uncertainty over the integrated representations before a choice is made. Task belief reflects the integrated representation that biases action, and task inference errors reflect the inference process following error feedback where the source of the task error is attributed to the response selected or input belief. In that work, we showed how these variables were encoded in frontoparietal cortices to facilitate integrative functions (Leach et al., 2025).

However, that work did not test how frontoparietal hubs coordinate information across distributed systems through connectivity. A key question, therefore, is how variables such as entropy and task inference errors modulate hub connectivity to regulate the convergence and redistribution of information across the brain. Addressing this question required a method that links trial-wise variables to adaptive changes in functional connectivity. Standard task-evoked functional connectivity approaches typically average across trials or use residual time series after removing task-evoked responses, limiting their ability to test whether trial-by-trial variables modulate network interactions (Cole et al., 2019; Di et al., 2021). To incorporate model-derived variables into connectivity estimation, we developed a model-based, task-related functional connectivity approach, based on beta-series and psychophysiological interaction approaches (beta-PPI). This method tests whether entropy, beliefs, and inference errors systematically modulate connectivity between frontoparietal hubs and distributed systems.

Applying this method to fMRI data, we examined how computational variables shape hub connectivity. Entropy increased connectivity between frontoparietal hubs and regions encoding task-relevant information. When uncertainty was high, hubs were more strongly coupled with regions representing inputs and outputs. Entropy also predicted fluctuations in task belief, which in turn modulated connectivity with action-related regions, suggesting a shift from integrating inputs to biasing outputs. Following feedback, task inference errors increased connectivity with input-related regions, consistent with updating beliefs. Together, these findings show how frontoparietal hubs implement integrative control by reconfiguring their connectivity through multiple computational signals, enabling flexible coordination of distributed brain systems across stages of integrative control.

## 2. Materials and methods

To test how frontoparietal hubs implement integration through their connectivity, we analyzed our previously published fMRI data (Leach et al., 2025). During the experiment, participants integrated an external sensory cue with an internal contextual state to determine the correct task on each trial (Fig. 1), allowing us to isolate distinct information streams and their integration. We developed a computational model to formalize the integration process (Fig. 2). The model quantified how sensory evidence and contextual information were integrated into a trial-wise joint probability distribution predicting the chosen task (task belief). It also generated entropy, reflecting uncertainty over the integrated inputs, and task inference error, defined as the discrepancy between the inferred task belief and the true task, which serves as a model-based updating signal following feedback. Together, these parameters provided trial-wise variables corresponding to input representation, integration, the resulting integrated task belief, and subsequent updating. In the present study, we extend our prior work by testing how these variables modulate the functional connectivity of frontoparietal connector hubs.

**Figure 1.**
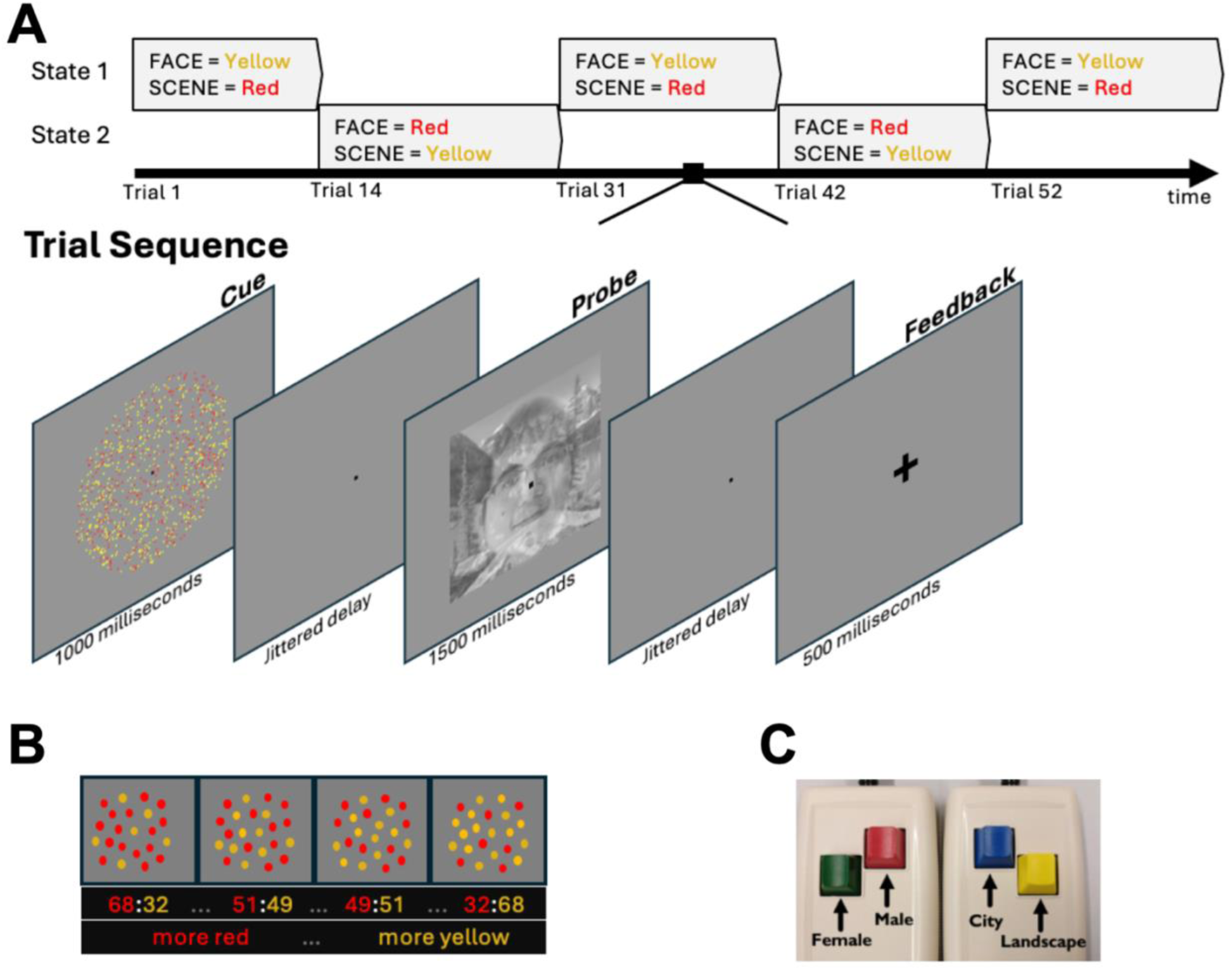
Behavioral Task. (A) Trial sequence and transitions of hidden state. Note the face image of this figure is the senior author of this paper, included only for illustration. (B) Color display. The dominant color was drawn from a continuous range between 0.51 and 0.68, thereby parametrically varying uncertainty in sensory evidence. (C) Response mappings.

**Figure 2.**
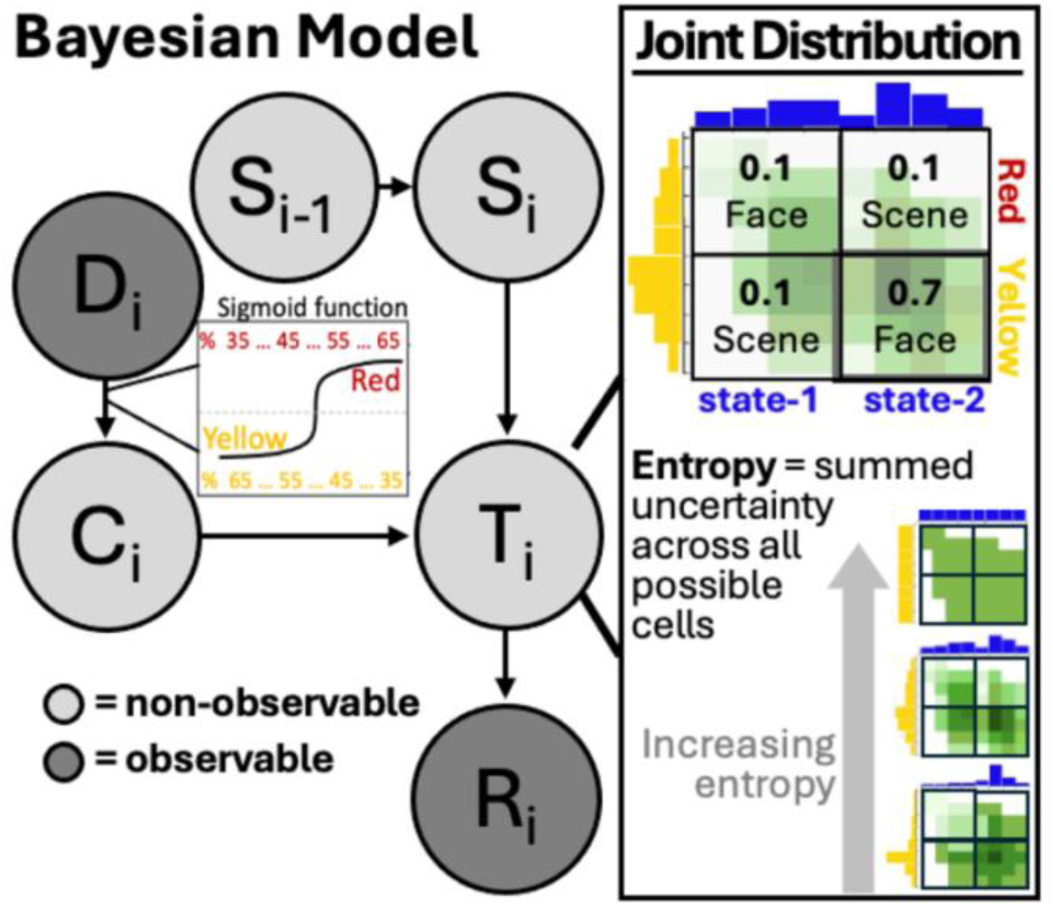
Computational model. Observable information (Ri and Di) was used to infer three latent variables on each trial i: state belief (Si), color belief (Ci), and task belief (Ti). A sigmoid function was applied to the presented color proportion (Di) to get an estimate of color belief (Ci). The state belief Si was inferred from Di, and Ri. Si and Ci were integrated into a 2×2 joint distribution from which the task belief Ti was inferred. The integration of Ci and Di produces uncertainty at the joint level. We quantified this integrated uncertainty using Shannon entropy of the 2×2 joint distribution. Entropy captures how uncertain the model is about the to-be-integrated inputs before a response is made.

### 2.1 Participants

We recruited 49 healthy participants (19 male and 30 female, *M*_age_ = 22.396, *SD*_age_ = 4.372, *Range*_age_ = 18-35 years) from the University of Iowa and the surrounding area to participate in this study. All participants had normal or corrected to normal visual acuity and color vision, were right-handed, and reported no history of epilepsy, psychiatric, or neurological conditions. This study was approved by the University of Iowa Institutional Review Board and conducted in accordance with the principles outlined in the Declaration of Helsinki. All participants gave written informed consent. Out of these 49 participants, 11 were excluded for excessive movement or chance-level behavioral performance during fMRI scanning, resulting in a final sample of 38 participants (15 male and 23 female, *M*_age_ = 22.95, *SD*_age_ = 5.88, *Range*_age_ = 18-35 years).

### 2.2 Behavioral task

To investigate integration, we developed a behavioral task that required subjects to integrate information from sensory inputs and an internally maintained state belief. Each trial began with a cue screen presented for one second. The cue consisted of an array of red and yellow dots moving randomly, and participants indicated whether the array contained more red or more yellow dots (Fig. 1A). The proportion of the dominant color was drawn from a continuous range between 51% and 68%, thereby parametrically varying uncertainty in sensory evidence (Fig. 1B). After a jittered delay (mean = 3.75 seconds; range = 0.5–10.5 seconds), a probe screen appeared for 1.5 seconds. The probe displayed a partially transparent face overlaid on a partially transparent scene image. On each trial, participants performed either a face task (male vs. female judgment) or a scene task (city vs. nature judgment). To correctly decide which task to perform, participants had to integrate perceptual color information from the cue with a hidden contextual state (Fig. 1A-B). In one hidden state, red signaled the face task and yellow signaled the scene task. In the other hidden state, these mappings were reversed: red signaled the scene task and yellow signaled the face task. The hidden state switched covertly every 10–20 trials. Response mappings were fixed across participants, with the left hand responded to the face task (index finger = male; middle finger = female), and the right hand responded to the scene task (index finger = city; middle finger = nature; Fig. 1C). Mappings were not counterbalanced. After a second jittered delay (mean = 3.75 seconds; range = 0.5–10.5 seconds), feedback was presented for 0.5 seconds. A plus sign indicated that the participant selected the correct task and made the correct perceptual discrimination. A minus sign indicated that the participant selected the incorrect task and/or made an incorrect discrimination, which signaled the need to infer the source of the error (i.e., whether the selected response was incorrect or whether the state or color belief was incorrect). Each trial ended with a jittered inter-trial interval (mean = 3.75 seconds; range = 0.5–10.5 seconds). Before scanning, participants completed a tutorial and practice session and were required to achieve at least 80% accuracy before proceeding. The scanning session consisted of five functional runs of 40 trials each (200 trials total). Each run lasted approximately 8.5 minutes.

### 2.3 Computational model

We developed a Bayesian model to formalize how participants integrated sensory evidence with contextual state information to determine which task to perform (Fig. 2). Full details and mathematical formulation of this computational model are reported in Leach et al. (2025) and also included in the Supplementary Materials (see also Supplementary Fig. 1). Here we describe the model conceptually, focusing on how the model provides trial-wise variables used in the subsequent analyses.

On each trial, the model estimates two probabilistic input representations, a state belief and a color belief. State belief represents the participant’s belief about the current hidden mapping (i.e., which color corresponds to which task). Because the mapping can switch unpredictably, this belief must be maintained and updated across trials using feedback. Color belief represents the participant’s belief about which color was dominant in the dot display. This belief reflects graded sensory evidence and perceptual uncertainty. These two probabilistic inputs are combined into a joint probability distribution that represents their integration. From this joint distribution, the model derives a task belief, a probabilistic estimate of which task (face or scene) should be performed on that trial (Fig 2, right panel). Following feedback, the model updates its beliefs using Bayes’ rule. We quantified updating using a task inference error, defined as the discrepancy between the model’s task belief and the true task outcome. Given that the task belief depends on the input beliefs, a larger task inference error indicates a significant mismatch between one input belief (either state or color) and its current true value. This signals a stronger need for the error inference process so that the source of this error (either color or state belief) can be properly attributed and updated.

Because both inputs are probabilistic, their integration produces uncertainty at the joint level. We quantified this integrated uncertainty using Shannon’s entropy of the 2×2 joint distribution. Entropy captures how uncertain the model is about the to-be-integrated inputs before a response is made. Low entropy corresponds to a sharply peaked joint distribution and high confidence in a particular state–color pairing. High entropy reflects a diffuse distribution and uncertainty across possible combinations (Fig. 2 bottom right panel). Entropy therefore provides a measure of integrated uncertainty during integration.

In our previous study (Leach et al., 2025), model comparisons showed that this model better predicts behavior than alternatives that do not integrate inputs through a joint probabilistic representation. For the present study, we extracted trial-wise estimates of state belief, color belief, task belief, entropy of the joint distribution, and task inference errors. State, color, and task beliefs were used for decoding analyses, whereas entropy, task belief, and task inference errors were used in connectivity analyses. Together, these variables allowed us to test frontoparietal hub connectivity patterns associated with input representations, integrated output, integrated uncertainty, and feedback-driven updating.

### 2.4 fMRI data acquisition and analyses

Imaging data were collected at the Magnetic Resonance Research Facility at the University of Iowa using a 3T GE SIGNA Premier scanner with a 48-channel head coil. Structural images were acquired using a multi-echo MPRAGE sequence (TR = 2348.52ms; TE = 2.968ms; flip angle = 8°; field of view = 256*256; 200 sagittal slices; voxel size = 1mm^3^). Functional images were acquired using an echo-planar sequence sensitive to blood oxygenated level-dependent (BOLD) contrast (multiband factor = 2; TR = 2039ms; TE = 30ms; flip angle = 75°; voxel size = 2 mm^3^).

#### 2.4.1 fMRI data preprocessing

All fMRI data were preprocessed using fMRIPrep version 23.2.0 (Esteban et al., 2019) to reduce noise and transform data from subject native space to the ICBM 152 Nonlinear Asymmetrical template version 2009c for group analysis (Fonov et al., 2009). Preprocessing steps include bias field correction, skull-stripping, co-registration between functional and structural images, tissue segmentation, motion correction, and spatial normalization to the standard space.

As part of preprocessing, we obtained nuisance regressors including rigid-body motion estimates (three translations and three rotations), cerebral spinal fluid, and white matter noise components from the component-based noise correction procedure. We also included derivatives and squares of these components. These nuisance regressors were entered in subsequent regression models to reduce the influences from noise and artifacts. We also censored the first three TRs of each functional run and any high motion TRs (framewise displacement greater than 0.5) to remove them from subsequent regression models. We did not perform any spatial smoothing when preprocessing the data.

#### 2.4.2 Parametric modulation analyses

We ran parametric modulation analyses to map brain regions where BOLD response amplitudes varied systematically with entropy. This was conducted GLMs using AFNI’s 3dDeconvolve with a voxel-wise restricted maximum likelihood estimate of a temporal correlation model using AFNI’s 3dREMLfit (Cox, 1996). Finally, we conducted mass univariate t-tests on the resulting beta values. Significant clusters for parametric modulation results are based on calculated multiple comparison corrected minimum cluster sizes of 128 voxels (cluster p-value threshold of 0.005 and a voxel p-value threshold of .05). This analysis allowed us to assess how entropy modulates the evoked-response magnitude beyond trial-averaged effects from the event epochs.

#### 2.4.3 Decoding analyses

The purpose of these analyses was to identify the neural substrates encoding (1) the input sources of information, state belief (S_i_) and color belief (C_i_) and (2) the output of this integrated product, task belief (T_i_). In short, we obtained trial-by-trial estimates of BOLD response amplitudes using a least-square-all approach implemented in AFNI’s 3dLSS. Decoding was performed via linear regression, with log-transformed trial-level model estimated participant beliefs (state belief, S_i_, color belief, C_i_, or task belief, T_i_) serving as the predicted variable, and the trial-level, voxel-wise beta estimates from 3dLSS as features. We applied a whole-brain searchlight decoding approach with an 8mm radius for each searchlight sphere. Within each searchlight sphere, voxel features were standardized by removing the mean and scaling to unit variance. For decoding, trial-wise betas were split into five groups based on the five functional runs, and a leave-one-group-out cross-validation approach was used to predict the trial-wise beliefs. Significant clusters for decoding results are based on calculated multiple comparison corrected minimum cluster sizes of 128 voxels (cluster p-value threshold of 0.005 and a voxel p-value threshold of .05).

### 2.5 Connector hub analyses

To map connector hubs, we calculated participation coefficient (PC) values, which measure how diversely connected a region is with distributed functional networks (Gratton et al., 2012). We calculated PC with two datasets. First, we ran these network hub analyses on an independent resting state dataset (Holmes et al., 2015) previously reported in Reber et al (2021) and Leach et al (2025). Second, we computed PC using task-residual data from the current dataset. Specifically, task-evoked responses modeling cue, probe, and epoch events were removed, and residual time series were used to estimate functional connectivity between 400 regions of interests (ROIs) defined by the Schaefer cortical parcellation (Schaefer et al., 2018).

Connectivity matrices were computed using pairwise Pearson correlation across ROIs, within each subject, and then averaged across subjects before calculating PC.

The PC value for each ROI i, PC*_i_* is defined as

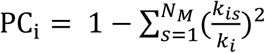

where *k_i_* is the sum of the total functional connectivity weight for ROI *i*, *k_is_* is the sum of the functional connectivity weight between ROI *i* and the cortical network *s*, and *N_M_* is the total number of cortical networks. To define cortical networks across ROIs, we used the 17 network parcellations from Yeo et al. (Yeo et al., 2011). These PC values ranged from 0 to 1, with a greater value indicating greater connectivity (i.e., greater “hub”-like properties). PC values were calculated across a range of network densities (.01 to .15) then averaged. To select hubs ROIs, we identified the top 20% of frontoparietal regions with the highest PC values (total 80 ROIs), using the independent resting-state dataset to avoid circularity between ROI definition and subsequent functional connectivity analyses.

### 2.6 Model-based functional connectivity

#### 2.6.1 Overview

To test how trial-wise computational variables (entropy, task belief, task inference errors) modulated functional connectivity between frontoparietal regions and distributed ROIs, we ordinally combined a beta-series connectivity approach (Rissman et al., 2004) with a generalized psychophysiological interaction approach (Friston et al., 1997; McLaren et al., 2012). This beta-PPI analysis consisted of three stages (Fig 3).

**Figure 3.**
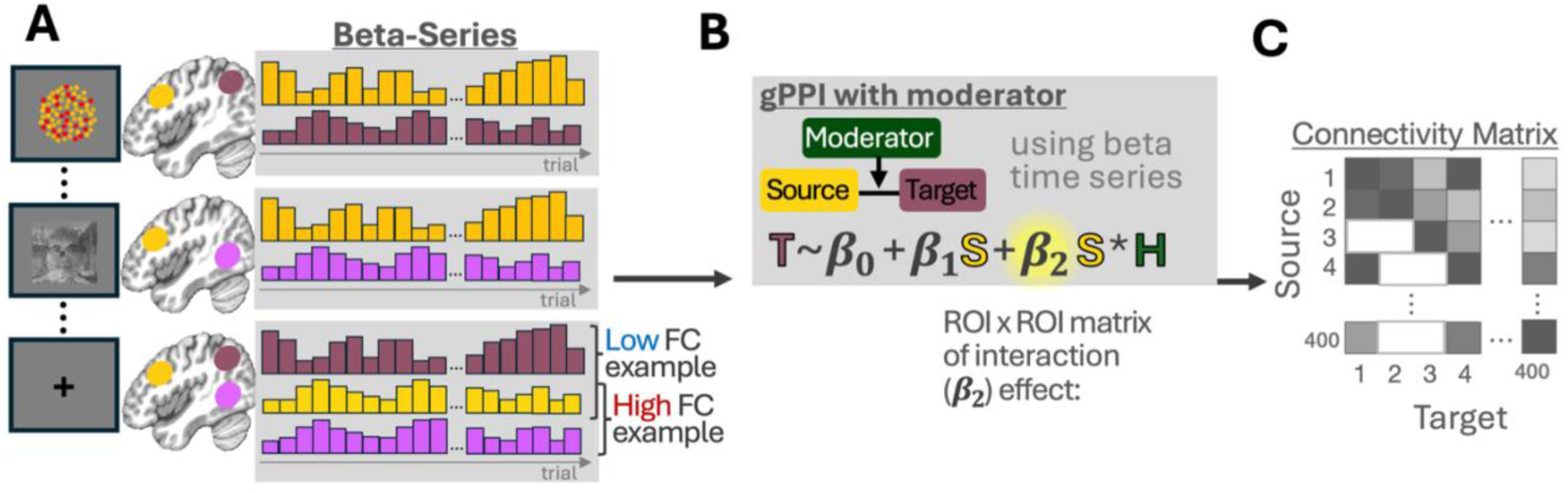
Task functional connectivity analysis. (A) Signel trial beta estimates were first obtained for each trial epoch and each ROI, resulting in timeseries of epoch specific trial-wise beta-series across ROIs. Note the face image of this figure is the senior author of this paper, included only for illustration. (B) A generalized PPI method was adapted to test how trial-wise variables (moderators) modulate functional connectivity between source and target regions. (C) Target by Source modulation matrices were submitted to a NBS procedure to identify significant connected edges within the matrix.

First, we estimated single-trial activity magnitudes (“beta-series”) for each event epoch (cue, probe, and feedback) using AFNI’s 3dLSS. Series of these single-trial, epoch-specific beta values provides a proxy for the trial-evoked activity fluctuations in each ROI (Fig 3A). Second, we used these beta series to estimate task-evoked connectivity by modeling relationships between ROIs using a psychophysiological interaction regression approach. Specifically, each model treated one ROI as the target (target ROI) and another as the source (source ROI). The regression included the source beta, the moderator (a trial-wise variable), and their interaction. The interaction term captured modulation of source–target functional connectivity by the trial-by-trial variable (Fig 3B). Third, we aggregated subject-level source-by-target interaction connectivity matrix to the group level and used a network-based statistic (NBS) procedure (Zalesky et al., 2010) to identify edges whose interaction effects were consistently different from zero across subjects (Fig 3C). Above steps were conducted separately for cue, probe, and feedback epochs, yield separate connectivity estimates for each trial epoch. Implementation details are described below.

A key motivation for this approach is that the model-derived variables of interest vary on a trial-by-trial basis. In addition, our experimental design imposes further constraints that limit the applicability of conventional task-based connectivity methods. Our task utilizes a randomized event-related design with multiple temporally adjacent epochs (cue, probe, feedback) separated by jittered intervals. In such designs, trial events are not grouped into discrete, repeating conditions, and the temporal overlap of hemodynamic responses makes it difficult to isolate condition-specific connectivity using standard PPI or gPPI approaches. Thus, standard functional connectivity methods that rely on residual time series or condition-averaged responses, with the assumption that the underlying modulation is stable across trials within the same condition, are not suitable for our study.

The beta-PPI approach addresses these limitations by separating estimation and interaction. First, single-trial responses are estimated independently for each event, reducing temporal overlap and allowing signals from adjacent trial epochs to be dissociated. Second, connectivity is modeled at the level of trial-wise responses, enabling continuous computational variables to be incorporated as modulators of connectivity. This framework therefore allows us to test whether connectivity between regions scales with internal computational states, rather than simply differing between predefined task conditions.

An additional advantage of this approach is that it reduces ambiguity in the interpretation of connectivity effects. The baseline term in the PPI model captures condition-independent coupling between regions, which can reflect shared task-evoked responses or co-activation rather than true interactions (Cole et al., 2019). In contrast, the interaction term in the PPI model isolates changes in coupling that scale with trial-wise variables, providing a more specific test of connectivity modulation. The trial-wise moderator is also included as a main-effect regressor, ensuring that variance related to evoked responses (including those modulated by the variable itself) is accounted for separately. As a result, the interaction term reflects connectivity changes beyond general activation or coactivation effects. By modeling trial-level responses and their interaction with trial-wise variables, beta-PPI separates general co-activation from task-relevant changes in inter-regional communication. The beta-PPI approach also preserves the temporal specificity of the computational model. Because model-derived variables vary continuously across trials, aligning them with single-trial neural responses avoids information loss from averaging across blocks of trials. Together, these features make the beta-series PPI framework well suited for our purpose of linking model-derived variables to task-related changes in functional connectivity in an event-related design. Below we describe its specific implementation.

#### 2.6.2 Beta-series

Single-trial beta estimates were obtained separately for each epoch (cue, probe, feedback) using AFNI’s least-squares–separate (LSS) deconvolution program (3dLSS). The final output consists of one beta estimate per voxel for each epoch and each trial. Separately for each epoch, beta series were then extracted from 400 cortical ROIs (Schaefer et al., 2018) and 17 subcortical regions of interests including the basal ganglia from the Harvard-Oxford subcortical atlas (Desikan et al., 2006), and different thalamic nuclei from the Morel atlas (Krauth et al., 2010).

These beta estimates reflect timeseries of cue, probe, or feedback activity that fluctuated across trials.

#### 2.6.3 PPI

Pairwise ROIs were then fitted with the following generalized psychophysiological interaction (gPPI) framework (McLaren et al., 2012) using ordinary least squares separately for each model-derived variable as moderator (T = target, S = source, H = moderator).

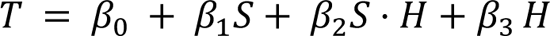

In contrast to conventional PPI approaches, which operate on continuous time series and require explicit hemodynamic convolution and deconvolution steps, our implementation was performed on trial-wise beta-series estimates derived from the beta-series procedure (see 2.6.2). Because each beta estimate already reflects the hemodynamically convolved BOLD response, no additional HRF convolution or deconvolution was applied at the PPI stage.

With this approach, we obtained a ROI by ROI matrix of β_1_ and β_2_. Please note β_3_ describes the effect of trial-wise variables on evoked responses, reported in our previous study and are not analyzed further in this study. Here, β_1_ indexes the subject’s baseline coupling of functional connectivity between those two regions, while the interaction term β_2_captures how trial-to-trial fluctuations in the variable (e.g., entropy, task belief, task inference errors) amplify or attenuate that coupling. This interaction matrix is the primary focus of the remaining analyses, and served as input for group NBS testing to determine significant task modulated functional connectivity (Zalesky et al., 2010). Each element within the matrix will be referred to as edges (connections between ROIs) hereon.

#### 2.6.4 Network based statistics

To identify reliable task modulation effects in functional connectivity at the group level, we applied the NBS approach (Zalesky et al., 2010). For each connection (edge) between regions, we first tested whether the interaction effect (β_2_) differed from zero across participants using a one-sample t-test. Rather than evaluating each edge independently, NBS identifies clusters of interconnected edges that show robust effects. Specifically, suprathreshold connections were grouped into connected components. These components capture clusters of edges whose connectivity is modulated by the model variable. For each component, we computed a summary statistic summing the overall strength of the effect across its constituent edges, and determine its statistical significance by comparing it to an empirically derived null distribution. To control for multiple comparisons, we generated a null distribution using cluster-based permutation testing.

On each permutation, the sign of each participant’s connectivity estimates was randomly flipped, and the full analysis was repeated to identify the largest component observed under the null hypothesis. This procedure yielded a distribution of maximal component sizes expected by chance. Observed components were assigned p-values based on this null distribution, and components with p ≤ 0.05 were retained. All connections belonging to these components were retained as showing reliable modulation of connectivity by the model variable. To facilitate interpretation, we summarized significant effects at the regional level by summing the overall strength of significant edges associated with each region.

#### 2.6.5 Summarizing connectivity estimates in independent ROIs

We further summarized significant edges for ROIs that showed significant decoding of source information (Section 2.5.2). Briefly, we first identified unbiased, independent ROIs with significant color, state, and task decoding by overlapping the Schaefer 400 atlas with cluster-corrected decoding maps. Independent ROIs were selected if more than 25% of their voxels showed above-chance decoding from the corrected decoding maps reported in Leach et al., 2025. For these ROIs, we then summed the weights of significant edges for each subject and compared them to the summed edge weights of ROIs that did not overlap with any significant decoding voxels using a within-subject paired t-test. These summaries provide a measure of how strongly each region participates in connectivity patterns modulated by the model variables.

#### 2.6.6 Singular value decomposition of connectivity matrix

To characterize the spatial structure of how task belief modulates connectivity, we performed a singular value decomposition of the group-averaged task belief interaction matrix (β_2_). The matrix was decomposed using singular value decomposition (SVD), yielding orthogonal components that capture dominant patterns of connectivity. Component loadings in brain space were projected back into atlas space using the same parcellation via inverse transformation. Variance explained by each component was quantified from the squared singular values, normalized by their sum.

#### 2.6.7 Simulation validation

To evaluate the sensitivity of the beta-PPI approach to detect modulation of connectivity at the BOLD level, we conducted a set of simulations that mirrored the full analysis pipeline.

These simulations were grounded in the task design and sample characteristics. Trial timing and were taken directly from the empirical dataset (38 subjects, 5 runs per subject), preserving the original timing distribution of cue, probe, and feedback events. Trial-wise modulatory variables (entropy) were also derived from the empirical data.

For each subject, latent neural signals were generated for a simulated source and target region. The source signal was constructed as a random vector of trial-wise task-related amplitudes. The target signal was then generated as a linear combination of the source signal, a task-independent modulatory term, and a source-by-modulator interaction term. Specifically, target_amp = β_1_ × source_amp + β_2_ × source_amp × entropy + β_2_ x entropy. For the simulation, β_1_ was set at fixed value of 0.38, which was the empirical median from the observed data. The strength of the interaction term served β_2_ as the primary effect-size parameter and was systematically varied across simulations. To ensure comparability across simulation conditions, the interaction coefficient was normalized to directly corresponds to a standardized effect size, such that a unit increase reflects a one standard deviation change in the target signal associated with a one standard deviation increase in the interaction term, holding other components constant.

These simulated neural signals were converted to BOLD time series using canonical hemodynamic response convolution of the gamma function. Measurement noise was then added to both source and target signals, with noise scaled to achieve a specified signal-to-noise ratio (SNR). Temporal autocorrelation was introduced to noise signals using an autoregressive process with parameters estimated from empirical residual data. Subject-level beta-series were estimated using the same LSS approach implemented via design matrices, matching the analysis pipeline applied to the real data. For each epoch (cue, probe, feedback), trial-wise source and target beta series were extracted and entered into the same PPI regression model in section 2.6.4. The interaction coefficient was tested using a two-sided t-test at the subject level.

To quantify sensitivity, simulations were repeated across a grid of interaction effect sizes and BOLD SNR values. For each parameter combination, 1000 simulations were performed. power was defined as the proportion of simulated samples in which the interaction effect was significant at a nominal alpha level of 0.05.

#### 2.6.8 Empirical SNR

To compare beta-PPI simulation results across SNR to empirical data, it is necessary to estimate the observed SNR. Empirical SNR was estimated from the LSS model by reconstructing the task-evoked BOLD signal and comparing it to residual variance. For each subject, we first extracted time series for the 400 ROIs from the GLM residual data and obtained trial-wise beta estimates from the LSS estimator for each task epoch (cue, probe, feedback). Using the original event onsets and durations, we then reconstructed the predicted BOLD signal by summing the contribution of each trial, computed as the product of its beta weight and the corresponding double gamma model. This yielded a model-predicted time series reflecting task-related signal.

Residual time series were calculated as the difference between the empirical signal and the task-related signal. Empirical SNR was then computed as the ratio of the standard deviation of the reconstructed signal to that of the residuals. Empirical SNR was computed separately for each ROI and each subject, and with the range of empirical SNR compared to the simulated SNR in section 2.6.5

### 2.7 Data and code availability

Code and data are available at https://github.com/HwangLabNeuroCogDynamics/FP-Hub-FC and https://openneuro.org/datasets/ds006821.

## 3. Results

The overarching goal of this project is to test how frontoparietal hub connectivity integrates information from diverse sources. We first verified that our beta-PPI approach can recover modulation of connectivity at the BOLD level using empirical simulations. We then examined whether frontoparietal regions exhibit connector hub properties in our task-based functional connectivity data, consistent with prior fMRI studies. Finally, we tested how trial-wise variables modulate the connectivity of frontoparietal hubs to support the integration of multiple sources of information.

### 3.1 Testing the sensitivity beta-PPI approach with Simulated Data

We first tested whether the beta-PPI approach can detect the modulation of connectivity at the BOLD level with our jittered task timing. To do this, we evaluated statistical power as a function of the interaction effect magnitude and BOLD signal-to-noise ratio (SNR) for each task epoch (cue, probe, feedback; Fig. 4). The overall statistical power was consistent across epochs, with similar profiles relating interaction effect size and SNR to detection probability. This indicates that sensitivity of the beta-series PPI approach was comparable across task epochs, despite differences in event timing and duration.

**Figure 4.**
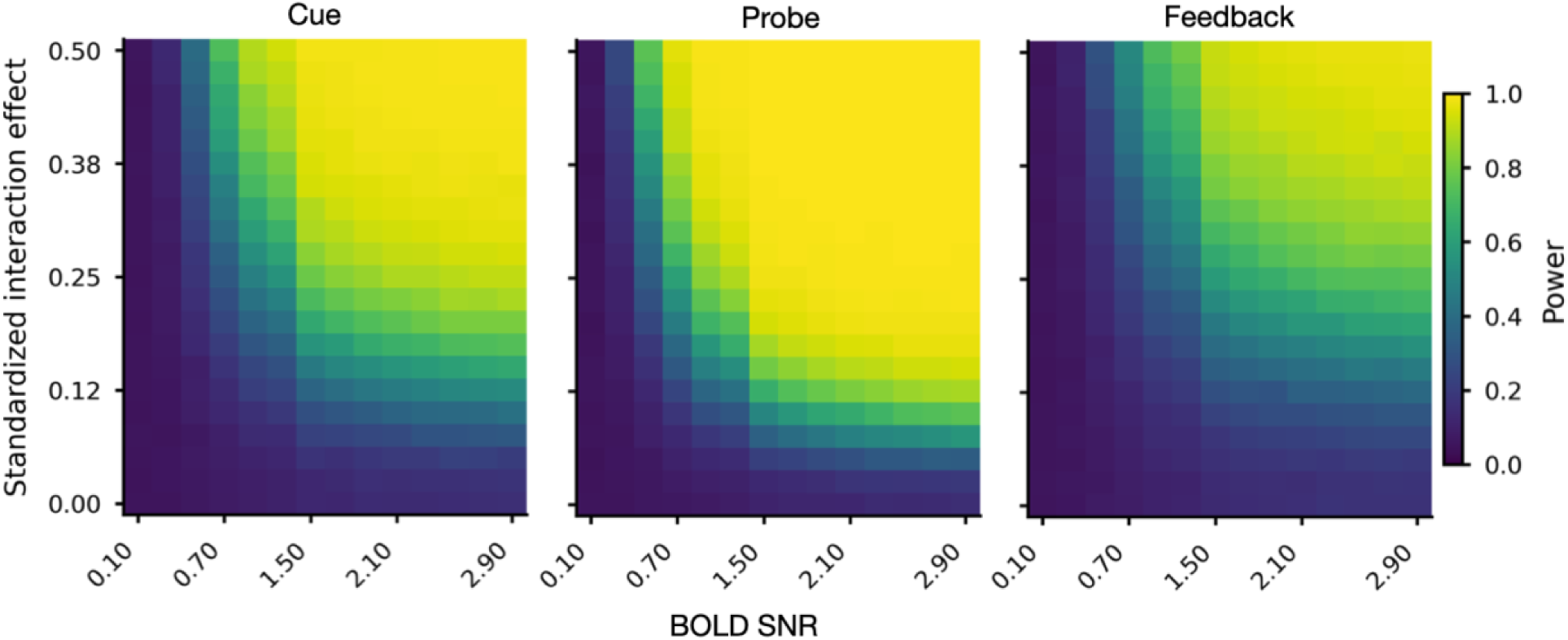
Beta-PPI simulation. Heatmaps show statistical power (two-sided, α = 0.05) for detecting the interaction term across simulated datasets, plotted as a function of standardized interaction effect (y-axis) and BOLD SNR (x-axis), separately for each task epoch.

Across all epochs, power increased monotonically with both interaction magnitude and SNR. Approximately 80% power was achieved when SNR exceeded ∼1.5 and the interaction parameter was in the range of 0.1 to 0.2. Notably, these values are comparable to the 80^th^ percentile interaction effects observed in the empirical data (β2 ≈ 0.13), and within the range of the observed BOLD SNR (25^th^ to 75^th^ percentile = 1.5 to 2.5), indicating that the observed effects are within the range associated with high statistical power under realistic SNR conditions.

### 3.2 Connector hubs

Having demonstrated that the beta-PPI approach can detect task-related functional connectivity, we next tested whether our data can detect the known connector hub architecture of the frontoparietal system (Bertolero et al., 2015; Gratton et al., 2012; Power et al., 2013). We quantified connector hub properties using the participation coefficient (PC) across two independent analyses. First, with an open access resting-state data analyzed in our previously published paper (Holmes et al., 2015; Reber et al., 2021), and second, with task-residual functional connectivity in the current dataset.

As shown in Fig. 5, frontoparietal regions exhibited strong connector hub properties across the two analyses. High PC regions were consistently localized to frontoparietal cortex, replicating prior reports of its role as a connector hub system in resting-state data (Bertolero et al., 2015; Cole et al., 2013), and extending this pattern to task-related connectivity in the current dataset. This convergence across independent datasets and analysis approaches indicates that the connector hub architecture is robustly expressed in our data and provides the foundation for us to further test how these hubs support integration through task-related modulation in functional connectivity. To ensure unbiased definition of hub regions and to avoid circularity, we selected frontoparietal ROIs based on PC values computed from the independent resting-state dataset, rather than the task-residual data, for subsequent analyses.

**Figure 5.**
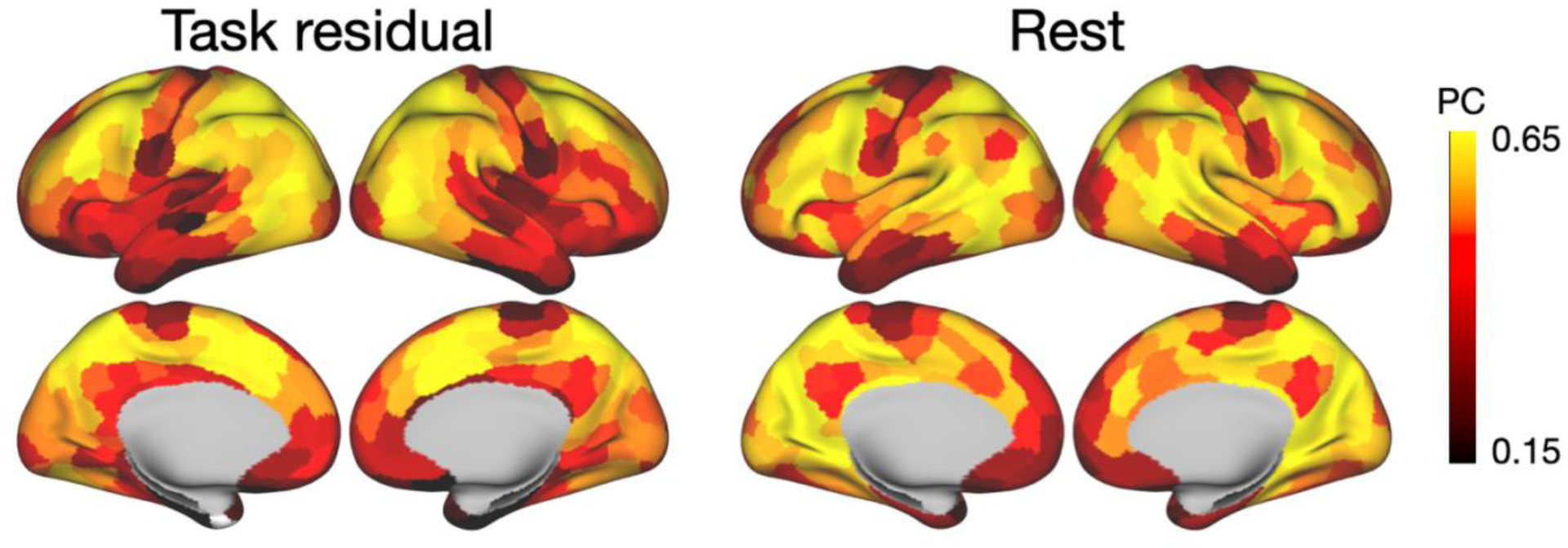
Connector hubs mapped in an independent resting-state data (right) and from task-residual functional connectivity of the current dataset (left).

### 3.3 Model-based connectivity analyses

Before presenting the connectivity results, we briefly summarize the model-derived variables used in the subsequent analyses. In our model, sensory evidence and contextual state are combined into a joint probability distribution that represents their integration on each trial (Fig 2). From this joint probability distribution, reflecting integrated representation, we derive three related but functionally distinct trial-wise variables that correspond to different stages of processing. Entropy quantifies uncertainty over the joint distribution prior to a decision, task belief reflects the inferred task derived from this distribution, and task inference errors index the magnitude of the feedback signal for error attribution (to state or color belief), driving input belief updating. Because all three variables are computed from the same integrated representation, a model-based functional connectivity analysis can test how integrated representations modulate connectivity between frontoparietal hubs and distributed brain regions for integrative control.

### 3.4 Entropy modulates frontoparietal functional connectivity with information sources

In our previous study (Leach et al., 2025), we found that entropy was behaviorally relevant, predicted trial-by-trial behavioral performance, with higher entropy predicting reaction time under conditions of increased uncertainty. From the fMRI parametric analysis, we also found that entropy increased evoked activity in frontoparietal regions that largely overlapped with areas exhibiting strong connector hub properties (Fig. 6A; spatial correlation across 417 ROIs = 0.38, p < 0.01). This pattern suggests that frontoparietal hubs encode integrated uncertainty across input sources and can potentially use this signal to regulate information transfer across distributed systems. Under conditions of high uncertainty, effective integration may require increased gain in both information representation and communication between frontoparietal hubs and regions encoding task-relevant inputs and outputs. Based on this hypothesis, we predicted that entropy would increase functional connectivity between frontoparietal hubs and distributed brain regions encoding distinct sources of information in the task.

**Figure 6.**
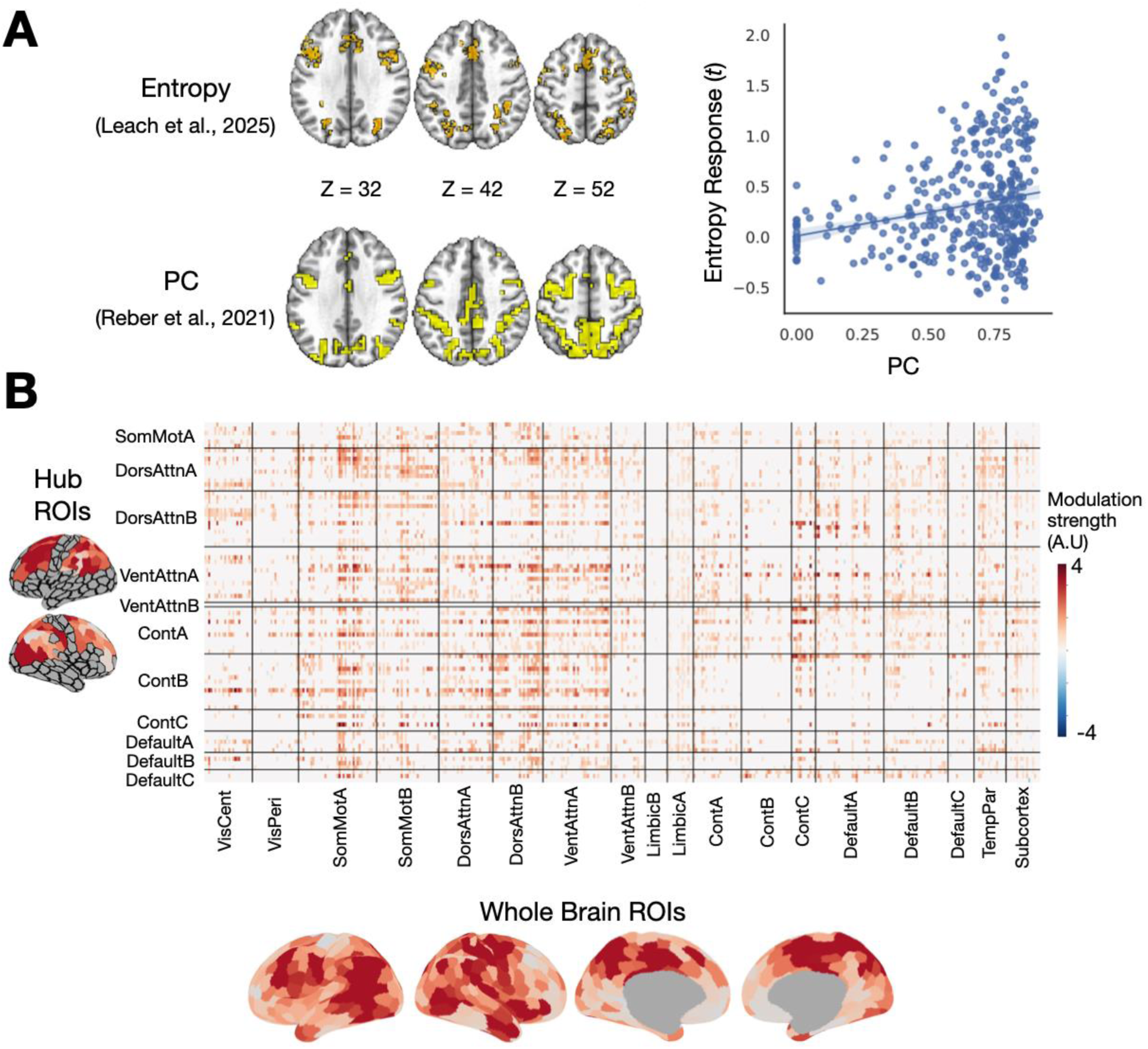
Entropy increased frontoparietal functional connectivity. (A) Left: frontoparietal regions show strong spatial correspondence between entropy-related modulation and connector hub properties (participation coefficient, PC). The entropy modulation map is from Leach et al. (2025), and the connector hub (PC) map is from Reber et al. (2020). Right: Scatter plot of entropy vs PC across ROIs. (B) Entropy positively modulates functional connectivity between frontoparietal hubs (rows of the connectivity matrix) and whole-brain ROIs (columns). Only significant edges that survived cluster correction at *p* < .05 corrected are displayed in the matrix. Rows and columns of the matrix were organized by 17 functional networks (SomMot: Somatomotor; DorsAttn: Dorsal Attention; VentAttn: Ventral Attention; Cont: Control; VisCent: Visual Central; VisPeri: Visual Peripheral; TempPar: Temporal Parietla). Surface maps (bottom) show the ROI-wise sum of entropy-related interaction coefficients across significant edges.

To test this prediction, we used the Schaefer 400-ROI atlas to identify top 20% regions in the frontoparietal cortices with strong connector hub properties, defined using PC values calculated from the independent resting-state dataset to ensure unbiased ROI selection (Fig. 6A; Holmes et al., 2015; Reber et al., 2021). We then examined how entropy modulated their functional connectivity with the rest of the brain. We focused on the cue epoch, when task-relevant information is first presented and must be integrated to guide subsequent behavior, making this period critical for testing how uncertainty influences information integration. We quantified entropy-related modulation of connectivity using the beta-PPI approach applied to trial-wise evoked activity. For each pair of ROIs, we designated one ROI as target and one as source, we then modeled trial-wise target activity as a function of source activity, entropy, and their interaction. The interaction term indexed the extent to which entropy modulated functional connectivity between regions. Critically, both the main effect of functional coupling and the entropy regressor were included in the model, such that any variance related to evoked responses is accounted for independently. As a result, the interaction term isolates connectivity changes that cannot be explained by overall activation differences, including those modulated by entropy. This strengthens the interpretation that the observed effects reflect entropy–related modulation on functional connectivity, rather than coactivation driven by general uncertainty. This analysis produced, for each subject, a matrix of entropy-dependent connectivity effects across all ROI pairs. At the group level, we applied a NBS permutation procedure to identify connected sets of edges showing reliable entropy-related modulation across subjects (Fig 6B). This approach controls for multiple comparisons while preserving the network structure of connectivity effects, allowing us to detect distributed patterns of entropy-modulated functional connectivity.

We found that entropy modulated functional connectivity between frontoparietal hubs and distributed regions across the brain (Fig. 6B, 19% of edges showed significant effects after correction), including bilateral motor regions, precentral sulcus, and medial frontal and parietal regions. This distribution was broad and spanned multiple functional networks used in the PC calculation, including the somatomotor, visual, default mode, and frontoparietal systems. Notably, the modulation effects were exclusively positive. This pattern indicates that entropy increased, rather than decreased, functional connectivity. Importantly, this effect was specific to entropy.

Other variables (task belief and task inference errors) tested did not yield significant connectivity modulation, indicating a selective role for entropy in regulating large-scale communication when multiple sources of inputs require integration.

To characterize which regions showed the strongest modulation, we identified independent cortical and subcortical ROIs that overlapped with our previously reported searchlight decoding results for task-relevant information (Fig. 7A). Color information was most strongly decoded from intraparietal, premotor, and lateral frontal regions; state information from rostral frontal, posterior parietal, and mediodorsal thalamic regions; and task information from bilateral primary motor, intraparietal, and caudal frontal regions (Leach et al., 2025). We then summarized the strength of entropy-related connectivity between frontoparietal hubs and these regions by summing interaction effects across edges. These values were compared to connectivity with ROIs that did not show significant decoding of task-relevant information using paired *t* tests (Fig. 7B). Entropy significantly increased connectivity between connector hubs and regions encoding color, state, and task information, with stronger effects than those observed for other comparison regions (color vs other: *t*(37) = 2.99, p = 0.004, task vs other: *t*(37) = 4.99, p <0.001, state vs other: *t*(37), p = 0.013). This pattern indicates that entropy preferentially enhances coupling between frontoparietal hubs and distributed regions that carry task-relevant inputs and outputs, consistent with a role in coordinating information integration across the network.

**Figure 7.**
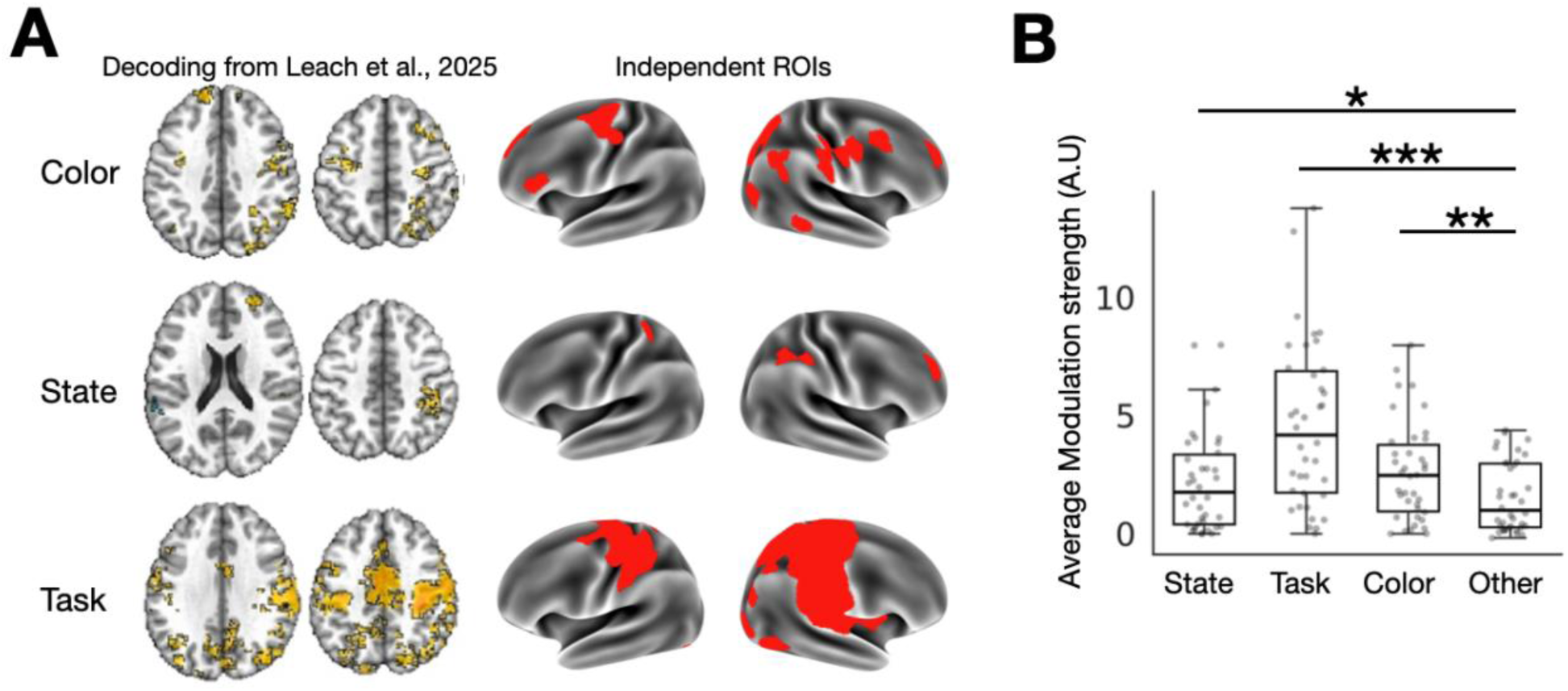
Entropy increased frontoparietal functional connectivity with ROIs encoding task-relevant inputs. (A) Independent ROIs (right) were selected based on searchlight decoding analyses (left; Leach et al., 2025) and included regions with >25% significant voxels encoding color, state, or task information. (B) Entropy preferentially enhances functional connectivity between frontoparietal hubs and regions encoding task-relevant inputs (color and state) and outputs (task), with stronger effects than in comparison ROIs that do not encode this information. Boxplots show the average of interaction coefficients for ROIs encoding color, state, and task information, compared with ROIs without significant decoding (other). * indicates p < 0.05; ** indicates p < 0.01; *** indicates p < 0.001.

### 3.5 Output task belief modulates frontoparietal functional connectivity with motor and occipitotemporal regions

After showing that entropy modulates frontoparietal connectivity during the cue epoch, we next examined how task belief influences connectivity during the probe epoch. Unlike the cue period, which requires integration of multiple information sources, subjects respond to the probe image base on the output of this integration process. In our computational model, task belief represents the inferred chosen task derived from the integrated joint distribution over inputs and therefore constitutes the integrated output guiding behavior at this stage.

We tested whether trial-wise task belief modulates functional connectivity between frontoparietal hubs and distributed brain regions using the same beta-PPI framework applied to probe-evoked activity. However, at the group level, we did not observe any connectivity clusters that survived correction for multiple comparisons for task belief or any other variables during the probe epoch. This suggests that, at the whole-brain level, connectivity modulation during this stage is weaker or more spatially focal than cue epoch integration.

However, our task design provides a strong a priori prediction for how task belief should modulate connectivity. In the experiment, task belief of the chosen task (face vs. scene) determines the corresponding motor effector (left vs. right hand; Fig 1C). In our model, task belief is parametrized as a scaler value from −1 to 1. Specifically, positive task belief values corresponded to the face task, whereas negative values corresponded to the scene task. This parameterization yields directional predictions, in which increasing (positive) task belief should be associated with stronger connectivity between frontoparietal hubs and the right motor regions controlling the left hand, while decreasing (negative) task belief should be associated with stronger connectivity with the left motor cortex. Because the task–effector mapping was not counterbalanced, it can serve as test of our integration model. Specifically, if task belief reflects the integrated output guiding behavior, it should modulate connectivity with the appropriate motor effector.

To assess whether the predicted patterns were present, we performed two exploratory analyses. We first examined the interaction effect matrix for the task belief interaction, by summing edge weights exceeding an uncorrected threshold for each ROI (p < 0.01; 2.4% of edges). Here we showed the results in axial slices of volume image, to better depict the expected motor effector regions (“hand knobs” for controlling hand movement)(Yousry et al., 1997). Positive task belief parameter, reflecting subjects chose the face task after integration, was associated with increased coupling between frontoparietal hubs and right motor cortex (Fig. 8A). In contrary, negative task belief, reflecting the scene task, showed the opposite pattern, with increased coupling to left motor cortex.

**Figure 8.**
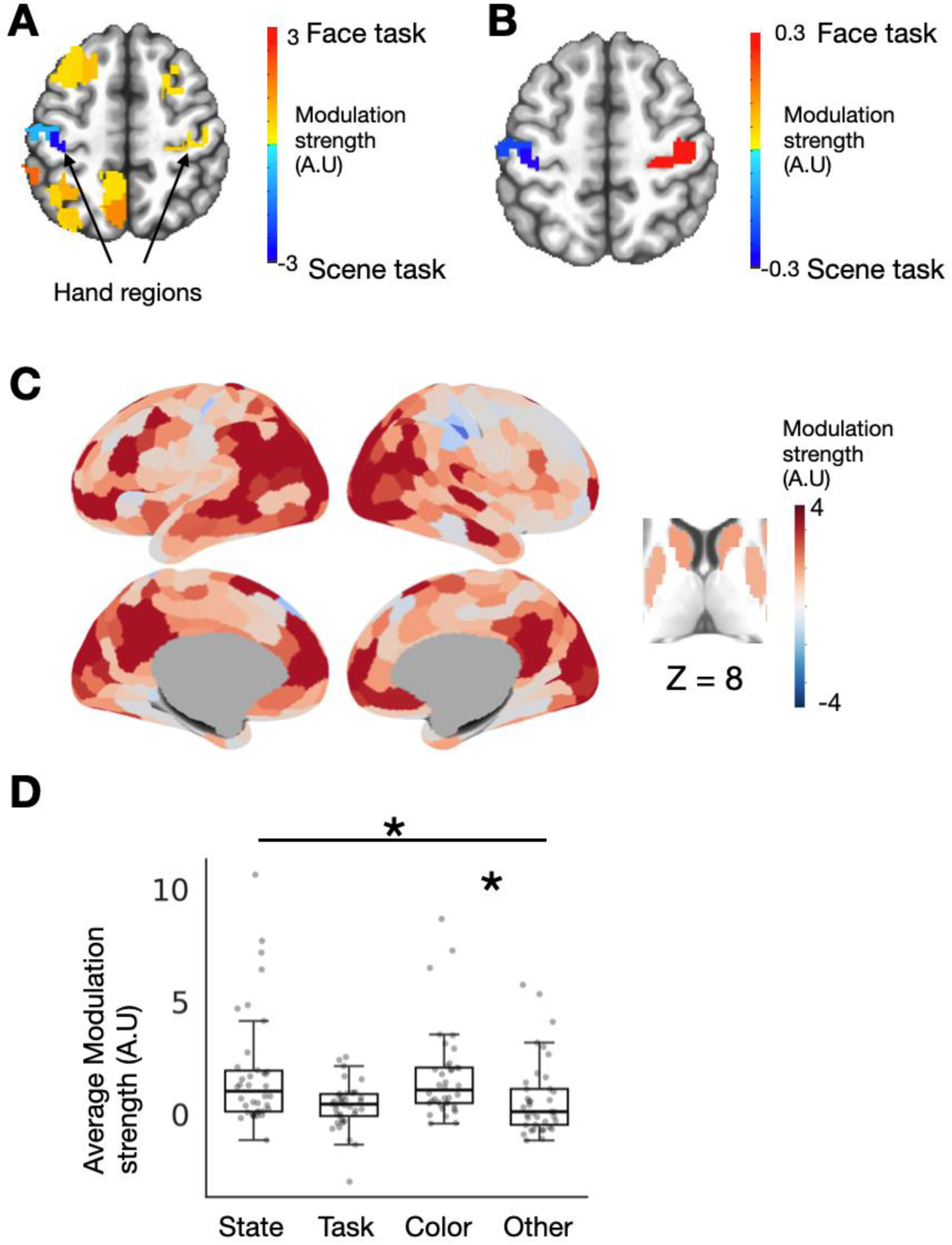
Task belief and task inference errors modulate frontoparietal functional connectivity during probe and feedback. (A) FC weights map showing task belief–related modulation of functional connectivity during the probe epoch. Positive task belief parameter (face task) is associated with increased coupling between frontoparietal hubs and the right motor cortex, whereas negative task belief parameter (scene task) shows increased coupling with the left motor cortex, consistent with the task structure. (B) Data-driven decomposition of the task belief–related connectivity pattern using singular value decomposition. The second-leading component revealed a lateralized organization, with stronger face-task belief associated with increased coupling between frontoparietal hubs and the right motor cortex, and stronger scene-task belief associated with increased coupling to the left motor cortex. The values reflect the loading weights from the component projected into atlas space. (C) Task inference errors modulate functional connectivity between frontoparietal hubs and whole-brain ROIs, including subcortical ROIs. The plot shows, for each ROI, the sum of interaction coefficients for inference errors-related modulation, after cluster-based correction at *p* < 0.05 corrected. (D) Task inference errors preferentially modulate connectivity between frontoparietal hubs and regions encoding task-relevant inputs. Boxplots show the averaged interaction coefficients for ROIs encoding color, state, and task information, compared to ROIs without significant decoding (other). * indicates p < 0.05; ** indicates p < 0.01; *** indicates p < 0.001.

To determine whether this spatial structure could be recovered in a data-driven manner, we then decomposed the interaction effect matrix for the task belief interaction term using singular value decomposition. This descriptive approach identifies low-dimensional patterns of functional connectivity modulated by task belief. One prominent component (Fig. 8B) exhibited a clear dissociation between tasks: stronger face-task belief was associated with increased coupling of frontoparietal hubs with the right motor cortex, whereas stronger scene-task belief showed increased coupling with the left motor cortex. This component accounted for 9.9% of the variance. Both analyses did not reveal specific components and activity patterns showing opposite directional modulation for the fusiform vs the parahippocampal areas. However, the resulting lateralized organization for the motor cortex closely matched the predicted effector-specific pattern, suggesting that frontoparietal hubs selectively modulate their connectivity with the motor systems in a manner consistent with the chosen task.

### 3.6 Task inference errors modulate frontoparietal functional connectivity during feedback

We next examined how model variables modulate frontoparietal connectivity during the feedback epoch. Unlike the cue and probe periods, which involve integration and task selection, the feedback epoch reflects updating based on feedback. In this context, task inference errors represent the discrepancy between the inferred task belief and the outcome and constitutes the final computational stage following entropy (uncertainty during integration) and task belief (the chosen task guiding behavior). Importantly, whereas entropy reflects uncertainty in task-relevant inputs prior to feedback, task inference error is a trial-specific signal that depends on the outcome and therefore captures a distinct updating process. As such, it provides a signal for updating internal representations. We therefore predicted that, during feedback, task inference error would modulate connectivity between frontoparietal hubs and distributed regions involved in representing task-relevant information. In contrast, we expected no significant modulation of connectivity by entropy or task belief during this epoch.

We tested this prediction using the beta-PPI framework applied to feedback-evoked activity. At the group level, only task inference errors showed significant modulation of functional connectivity during feedback. Entropy and task belief were also tested during the feedback epoch but did not show any significant modulation of connectivity, indicating that the observed effects are not driven by general uncertainty. As shown in Fig. 8C, task inference errors modulated connectivity between frontoparietal hubs and a widely distributed set of regions across the brain. Similar to entropy, these effects were spatially extensive, indicating that PE influences large-scale communication within the network. Notably, the direction of modulation varied across regions. Task inference errors was associated with increased connectivity between frontoparietal hubs and distributed cortical regions, including temporal other frontoparietal areas, and subcortical regions including the caudate and putamen, while showing reduced connectivity with motor cortex. This pattern suggests that larger PEs are associated with a decoupling of motor-related processing and an increase in coupling with regions involved in representing and updating task-relevant information. Of note, these effects reflect modulation captured by the interaction term in the beta-PPI model, indicating changes in connectivity rather than baseline coactivation.

To further characterize these effects, we performed the same region-of-interest analysis used for entropy by leveraging independently defined ROIs that encode task-relevant information (Fig. 8D). We summarized the strength of task inference errors related connectivity between frontoparietal hubs and these regions by summing interaction effects across edges and compared them to connectivity with ROIs that did not encode task-relevant information. Task inference errors significantly increased connectivity between frontoparietal hubs and regions encoding color and state information, but not regions encoding task belief (task vs. other: t(37) = −0.98, p > 0.05; state vs. other: t(37) = 2.27, p = 0.026; color vs. other: t(37) = 2.31, p = 0.023). This pattern indicates that inference errors preferentially enhance coupling with regions representing input features of the task, as well as updating internal state belief, suggesting a role in updating representations of the environment before integration to derive the output task rule.

Together, these results show that task inference errors selectively modulate large-scale functional connectivity during feedback, shifting coupling away from motor output systems and toward distributed regions encoding task-relevant inputs. The absence of entropy effects during this stage, combined with the specificity of the task inference errors interaction term, indicates that this modulation reflects updating processes rather than general uncertainty. This pattern complements the effects observed for entropy during the cue epoch and task belief during the probe epoch, suggesting that frontoparietal hubs reconfigure their connectivity to support distinct computational demands across integration stages.

## 4. Discussion

Frontoparietal cortex has long been implicated in the integration of information to support flexible, goal-directed behavior. Early theoretical accounts emphasized the role of transmodal association cortex in linking perception, memory, and action (Mesulam, 1998). These regions were proposed to sit at the apex of cortical hierarchies, enabling the convergence of diverse information streams (Damasio, 1989). More recent work has formalized this idea using network science approaches, quantifying frontoparietal regions as “connector hubs” that exhibit broad, distributed connectivity across multiple functional systems (Bertolero et al., 2015; Gratton et al., 2012; Power et al., 2013). Measures such as participation coefficient, network density, and articulation points have consistently highlighted frontoparietal cortices’ broad access to multiple functional networks. Together, these findings have established the topological basis for the hypothesis that frontoparietal hubs support integrative control.

A limitation of that evidence is that integration has largely been inferred from static measures of connectivity. Most studies rely on resting-state functional connectivity or structural connectivity derived from tractography, which characterize the organization of network architecture but do not directly measure how information is actively processed during behavior (Bassett and Sporns, 2017). Specifically, they did not investigate the computational processes that occur within hubs, nor how these processes shape communication with other brain regions.

To address this limitation requires advances on two challenges. First, to characterize the representations and computations implemented within frontoparietal hubs during integration. Second, to determine how these computational variables influence large-scale network interactions. In our previous work (Leach et al., 2025), we addressed the first challenge by developing a Bayesian model that formalizes how sensory evidence and contextual state information are integrated into a joint representation to achieve integration. This model yielded several model-derived variables, including joint probabilistic representations of multiple inputs, an integrated task belief, and entropy as a measure of uncertainty over the joint distribution. It is important to note that this model is distinct from multi-sensory integration, as it combines sensory inputs with internal representations, and thus are consistent with recent proposal of conjunctive representation, in which multiple task-relevant features can bridge inputs with adaptive actions (Ito et al., 2022; Kikumoto and Mayr, 2020). In addition to showing how frontoparietal representations jointly encode multiple inputs and integrated outputs, we further introduced entropy as a control-relevant signal that quantifies uncertainty over integrated representations. Therefore, entropy may signal when integration is uncertain and thus require increased coordination across distributed systems.

The present study addresses the second challenge, how trial-wise variables shape frontoparietal hubs connectivity with the rest of the brain. Using a model-based, trial-wise functional connectivity approach, we show that entropy, task belief, and task inference errors selectively modulate frontoparietal hub connectivity across different stages of behavior. At the stage of information presentation (the cue epoch), entropy increased functional coupling between frontoparietal hubs and regions encoding task-relevant inputs and outputs. This pattern suggests that uncertainty acts as a control signal that amplifies communication between hubs and the distributed systems that provide information for integration. Thus, our finding provide evidence from network level interactions on classic connectionist models that cognitive control increases the gain of task-relevant processing under conditions of uncertainty or conflict (Cohen et al., 1990).

During the probe epoch and after integrating sensory and state inputs, task belief selectively modulated connectivity between frontoparietal hubs and motor regions that are aligned with the chosen task. Because in our model, task belief is derived from the same joint distribution as entropy, it reflects the outcome of the integration process. The observed connectivity pattern suggests that frontoparietal hubs modulate their coupling with the motor systems based on the outcome of integration, consistent with top-down control over information processing (Miller and D’Esposito, 2005). Importantly, this effect was anatomically specific and constrained by the task structure (e.g., right motor cortex for the face task, left motor cortex for the scene task), indicating that connectivity modulation can selectively target relevant effector systems. Finally, following feedback, task inference errors modulated frontoparietal connectivity by increasing coupling with regions encoding task-relevant inputs and internal state, while reducing coupling with motor regions. Task inference errors was computed relative to task belief and therefore reflects the need to update internal representations when expectations are violated.

Taken together, these results show that entropy, task belief, and task inference errors act as distinct, stage-specific control signals that selectively modulate frontoparietal hub connectivity across different stages of integrative control. Although these signals exert dissociable effects on network interactions with each engaging different systems at different task epochs, they are all derived from a common integrative representation, the joint probability distribution that integrates task-relevant inputs (Leach et al., 2025). Within this framework, the joint distribution serves as a shared representation from which multiple control signals are read out at different stages of processing. Before a decision is made, entropy is computed over this distribution, capturing the degree of uncertainty across possible input combinations. From the same distribution, task belief is derived as the marginal and selected output, representing the most probable task given the integrated inputs. Following feedback, task inference errors is computed by comparing this inferred task belief to the observed outcome, quantifying the need to update beliefs prior to integration on subsequent trials.

This relationship suggests that frontoparietal hubs can potentially transform an integrated representation into multiple control signals that are deployed in a temporally and functionally specific manner. In this sense, hub connectivity may multiplex these signals to generate distinct patterns of network connectivity depending on current task demands. Such multiplexing allows the same underlying integration process to flexibly shift between enhancing communication with distributed input systems under uncertainty, biasing output pathways during decision, and re-engaging input systems during updating.

A key limitation of the currently study is that our methods do not establish the directionality of information flow between frontoparietal hubs and other brain regions. Although our analyses designate “source” and “target” regions within the beta-PPI method, this distinction reflects a statistical modeling terminology rather than a causal relationship. The interaction term captures how activity in one region covaries with another as a function of a model-derived variable, but it does not indicate whether frontoparietal hubs drive changes in downstream regions, receive input from them, or participate in bidirectional exchange (albeit the most likely scenario). As such, the observed modulation of connectivity should be interpreted as changes in functional coupling rather than directed causal connectivity. Future work using methods that explicitly model effective connectivity can resolve this limitation. Dynamic causal modeling, for example, provides a method for testing directed interactions and how they are modulated by task variables, and has been successfully applied to characterize cortical dynamics within the frontal cortex (Nee and D’Esposito, 2016). In addition, causal perturbation approaches such as transcranial magnetic stimulation or other related methods can test whether frontoparietal hubs exert a causal influence on distributed network functions (Hwang et al., 2020; Iyer et al., 2022; Nee and D’Esposito, 2017).

A second limitation is that the present analyses characterize group-level connectivity patterns, which may obscure meaningful individual differences in network organization (Gordon et al., 2017). Recent precision mapping studies have demonstrated that regions identified as connector hubs at the group level often correspond to areas of high network density, but there are also individual differences in high density hub zones that cannot be sensitively detected by group averaging approaches (Ladwig et al., 2025). Future studies leveraging high-resolution, within-subject mapping will be important for determining whether the integration effects observed here are implemented by distinct, individual specific hub regions.

In summary, the principal contribution of this study is to show how frontoparietal connector hubs implement integrative control by transforming an integrated probabilistic representation into multiple control signals that selectively reconfigure frontoparietal hub connectivity. This identifies a network mechanism by which integration is enacted, linking representations and control signals with network interactions across distributed brain systems.

## Acknowledgments

Author contributions are as follows.

Conceptualization: KH, SCL, JJ

Methodology: KH, SCL, JJ

Investigation: KH, SCL, SES, JJ

Visualization: KH, SCL, SES, JJ

Supervision: KH, JJ

Writing—original draft: KH, SCL

Writing—review & editing: KH, SCL, JJ

Authors declare that they have no competing interest.

All data and code will be made available upon publication.

Research reported here was supported by a National Institutes of Mental Health grants R01MH122613, R01MH140248, and the Iowa Neuroscience Institute. This work was conducted on an MRI instrument funded by S10OD025025.

## References

Bassett, D.S., Sporns, O., 2017. Network neuroscience. Nat. Neurosci. 20, 353–364.

Bertolero, M.A., Yeo, B.T.T., Bassett, D.S., D’Esposito, M., 2018. A mechanistic model of connector hubs, modularity and cognition. Nat Hum Behav 2, 765–777.

Bertolero, M.A., Yeo, B.T.T., D’Esposito, M., 2015. The modular and integrative functional architecture of the human brain. Proc. Natl. Acad. Sci. U. S. A. 112, E6798–807.

Cohen, J.D., Dunbar, K., McClelland, J.L., 1990. On the control of automatic processes: a parallel distributed processing account of the Stroop effect. Psychol. Rev. 97, 332–361.

Cole, M.W., Ito, T., Schultz, D., Mill, R., Chen, R., Cocuzza, C., 2019. Task activations produce spurious but systematic inflation of task functional connectivity estimates. Neuroimage 189, 1–18.

Cole, M.W., Reynolds, J.R., Power, J.D., Repovs, G., Anticevic, A., Braver, T.S., 2013. Multi-task connectivity reveals flexible hubs for adaptive task control. Nat. Neurosci. 16, 1348–1355.

Cox, R.W., 1996. AFNI: software for analysis and visualization of functional magnetic resonance neuroimages. Comput. Biomed. Res. 29, 162–173.

Damasio, A.R., 1989. The Brain Binds Entities and Events by Multiregional Activation from Convergence Zones. Neural Comput. 1, 123–132.

Desikan, R.S., Ségonne, F., Fischl, B., Quinn, B.T., Dickerson, B.C., Blacker, D., Buckner, R.L., Dale, A.M., Maguire, R.P., Hyman, B.T., Albert, M.S., Killiany, R.J., 2006. An automated labeling system for subdividing the human cerebral cortex on MRI scans into gyral based regions of interest. Neuroimage 31, 968–980.

Di, X., Zhang, Z., Biswal, B.B., 2021. Understanding psychophysiological interaction and its relations to beta series correlation. Brain Imaging Behav. 15, 958–973.

Esteban, O., Markiewicz, C.J., Blair, R.W., Moodie, C.A., Isik, A.I., Erramuzpe, A., Kent, J.D., Goncalves, M., DuPre, E., Snyder, M., Oya, H., Ghosh, S.S., Wright, J., Durnez, J., Poldrack, R.A., Gorgolewski, K.J., 2019. fMRIPrep: a robust preprocessing pipeline for functional MRI. Nat. Methods 16, 111–116.

Fonov, V.S., Evans, A.C., McKinstry, R.C., Almli, C.R., Collins, D.L., 2009. Unbiased nonlinear average age-appropriate brain templates from birth to adulthood. Neuroimage Supplement 1, S102.

Friston, K.J., Buechel, C., Fink, G.R., Morris, J., Rolls, E., Dolan, R.J., 1997. Psychophysiological and modulatory interactions in neuroimaging. Neuroimage 6, 218–229.

Fuster, J.M., 2015. The Prefrontal Cortex. Academic Press.

Goldman-Rakic, P.S., 1988. Topography of cognition: parallel distributed networks in primate association cortex. Annu. Rev. Neurosci. 11, 137–156.

Gordon, E.M., Laumann, T.O., Gilmore, A.W., Newbold, D.J., Greene, D.J., Berg, J.J., Ortega, M., Hoyt-Drazen, C., Gratton, C., Sun, H., Hampton, J.M., Coalson, R.S., Nguyen, A.L., McDermott, K.B., Shimony, J.S., Snyder, A.Z., Schlaggar, B.L., Petersen, S.E., Nelson, S.M., Dosenbach, N.U.F., 2017. Precision Functional Mapping of Individual Human Brains. Neuron 95, 791–807.e7.

Gratton, C., Nomura, E.M., Pérez, F., D’Esposito, M., 2012. Focal brain lesions to critical locations cause widespread disruption of the modular organization of the brain. J. Cogn. Neurosci. 24, 1275–1285.

Holmes, A.J., Hollinshead, M.O., O’Keefe, T.M., Petrov, V.I., Fariello, G.R., Wald, L.L., Fischl, B., Rosen, B.R., Mair, R.W., Roffman, J.L., Smoller, J.W., Buckner, R.L., 2015. Brain Genomics Superstruct Project initial data release with structural, functional, and behavioral measures. Sci Data 2, 150031.

Hwang, K., Shine, J.M., Cellier, D., D’Esposito, M., 2020. The Human Intraparietal Sulcus Modulates Task-Evoked Functional Connectivity. Cereb. Cortex 30, 875–887.

Ito, T., Yang, G.R., Laurent, P., Schultz, D.H., Cole, M.W., 2022. Constructing neural network models from brain data reveals representational transformations linked to adaptive behavior. Nat. Commun. 13, 1–16.

Iyer, K.K., Hwang, K., Hearne, L.J., Muller, E., D’Esposito, M., Shine, J.M., Cocchi, L., 2022. Focal neural perturbations reshape low-dimensional trajectories of brain activity supporting cognitive performance. Nat. Commun. 13, 4.

Kanwisher, N., Yovel, G., 2006. The fusiform face area: a cortical region specialized for the perception of faces. Philos. Trans. R. Soc. Lond. B Biol. Sci. 361, 2109–2128.

Kikumoto, A., Mayr, U., 2020. Conjunctive representations that integrate stimuli, responses, and rules are critical for action selection. Proc. Natl. Acad. Sci. U. S. A. 117, 10603–10608.

Krauth, A., Blanc, R., Poveda, A., Jeanmonod, D., Morel, A., Székely, G., 2010. A mean three-dimensional atlas of the human thalamus: generation from multiple histological data. Neuroimage 49, 2053–2062.

Ladwig, Z., Kermani, K.Z., Dworetsky, A., Labora, N., Hernandez, J.J., Dorn, M., Smith, D.M., Nee, D.E., Petersen, S.E., Braga, R.M., Gratton, C., 2025. Precision fMRI reveals densely interdigitated network patches with conserved motifs in the lateral prefrontal cortex. bioRxivorg. 10.1101/2025.07.24.666468

Leach, S.C., Hollow, H., Jiang, J., Hwang, K., 2025. Frontoparietal hubs leverage probabilistic representations and integrated uncertainty to guide cognitive flexibility. J. Neurosci. e0989252025.

McLaren, D.G., Ries, M.L., Xu, G., Johnson, S.C., 2012. A generalized form of context-dependent psychophysiological interactions (gPPI): a comparison to standard approaches. Neuroimage 61, 1277–1286.

Mesulam, M.M., 1998. From sensation to cognition. Brain 121 ( Pt 6), 1013–1052.

Mesulam, M.-M., 1990. Large-scale neurocognitive networks and distributed processing for attention, language, and memory. Annals of Neurology: Official Journal of the American Neurological Association and the Child Neurology Society 28, 597–613.

Miller, B.T., D’Esposito, M., 2005. Searching for “the Top” in Top-Down Control. Neuron 48, 535–538.

Nee, D.E., D’Esposito, M., 2017. Causal evidence for lateral prefrontal cortex dynamics supporting cognitive control. Elife 6. 10.7554/eLife.28040

Nee, D.E., D’Esposito, M., 2016. The hierarchical organization of the lateral prefrontal cortex. Elife 5. 10.7554/eLife.12112

Power, J.D., Schlaggar, B.L., Lessov-Schlaggar, C.N., Petersen, S.E., 2013. Evidence for hubs in human functional brain networks. Neuron 79, 798–813.

Reber, J., Hwang, K., Bowren, M., Bruss, J., Mukherjee, P., Tranel, D., Boes, A.D., 2021. Cognitive impairment after focal brain lesions is better predicted by damage to structural than functional network hubs. Proc. Natl. Acad. Sci. U. S. A. 118. 10.1073/pnas.2018784118

Rissman, J., Gazzaley, A., D’Esposito, M., 2004. Measuring functional connectivity during distinct stages of a cognitive task. Neuroimage 23, 752–763.

Schaefer, A., Kong, R., Gordon, E.M., Laumann, T.O., Zuo, X.-N., Holmes, A.J., Eickhoff, S.B., Yeo, B.T.T., 2018. Local-Global Parcellation of the Human Cerebral Cortex from Intrinsic Functional Connectivity MRI. Cereb. Cortex 28, 3095–3114.

Sporns, O., Betzel, R.F., 2016. Modular Brain Networks. Annu. Rev. Psychol. 67, 613–640.

Tononi, G., Sporns, O., Edelman, G.M., 1994. A measure for brain complexity: relating functional segregation and integration in the nervous system. Proceedings of the National Academy of Sciences 91, 5033–5037.

van den Heuvel, M.P., Sporns, O., 2013. Network hubs in the human brain. Trends Cogn. Sci. 17, 683–696.

Yeo, B.T.T., Krienen, F.M., Sepulcre, J., Sabuncu, M.R., Lashkari, D., Hollinshead, M., Roffman, J.L., Smoller, J.W., Zöllei, L., Polimeni, J.R., Fischl, B., Liu, H., Buckner, R.L., 2011. The organization of the human cerebral cortex estimated by intrinsic functional connectivity. J. Neurophysiol. 106, 1125–1165.

Yousry, T.A., Schmid, U.D., Alkadhi, H., Schmidt, D., Peraud, A., Buettner, A., Winkler, P., 1997. Localization of the motor hand area to a knob on the precentral gyrus. A new landmark. Brain 120 ( Pt 1), 141–157.

Zalesky, A., Fornito, A., Bullmore, E.T., 2010. Network-based statistic: identifying differences in brain networks. Neuroimage 53, 1197–1207.

